# Abnormally low expression of tumor necrosis factor stimulated gene 6 (TSG-6) in obese type 2 diabetic mice contributes to altered inflammation and delayed cutaneous wound healing

**DOI:** 10.64898/2026.09.19.752893

**Authors:** Minou Alipour, Joseph Offenberger, Sanjay Anand, Yan Wang, Olga Stenina-Adognravi, Edward V. Maytin

## Abstract

Type 2 diabetes mellitus (T2DM) is a chronic disorder associated with obesity and hyperglycemia. T2DM is linked to significant clinical complications such as delayed wound healing and diabetic foot ulcers. Tumor necrosis factor-stimulated gene 6 (TSG-6) is a protein with multiple functions including regulation of cytokine binding to glycosaminoglycans in extracellular matrices and on vascular endothelial walls. Loss of TSG-6 in knockout mice increases skin inflammation and delays wound healing. Here, in both a genetic model (*db/db* mice) and a dietary model that simulates human T2DM (mice treated with a high-fat diet [HFD] and streptozotocin [STZ]), we observed sustained hyperglycemia and a reproducible delay in wound closure. TSG-6 expression was markedly reduced in diabetic skin in these models, both before injury and post wounding. HFD+STZ diabetic mouse wounds showed elevated levels of pro-inflammatory cytokines (IL-6, TNF-α, and MIP-1α) and lower levels of the anti-inflammatory cytokine IL-10, relative to normoglycemic mice. Re-introduction of TSG-6 protein into the wounds normalized healing rates, corrected abnormal cytokine expression levels, and increased macrophage accumulation in dermal and adipose compartments. Thus, T2DM is associated with reduced levels of TSG-6 in diabetic skin, which contributes to the pro-inflammatory milieu and the delay in wound closure.

## INTRODUCTION

Type 2 diabetes mellitus (T2DM) is a subtype of diabetes caused by multiple genetic and environmental factors that result in pancreatic β-cell failure and increased resistance of peripheral tissues to the effects of insulin (Donath, 2019, Donath and Shoelson, 2011). Clinical complications of diabetes can involve the heart, kidneys, vasculature, peripheral nerves, and skin (Barrett et al., 2017, Gordois et al., 2003, Hoffstad et al., 2015). Of the estimated 500+ million people worldwide with diabetes, 20-30% will develop a non-healing wound (usually a diabetic foot ulcer), and amongst the latter, 20% will require a lower-extremity amputation (McDermott et al., 2023). In the U.S., overall healthcare costs for treating diabetes is >240 billion $/year), with 30% of that related to diabetic foot ulcer care (Waibel et al., 2023); the latter actually exceeds the cost for treating cancer (Armstrong et al., 2020).

Hyperglycemia is the major driver of diabetes and contributes to impaired wound healing through diverse mechanisms, including: (1) defective cellular immune responses (Berbudi et al., 2020), (2) impaired fibroblast responses (Liu et al., 2022), (3) increased oxidative stress (Giacco and Brownlee, 2010), (4) microvascular damage and delayed angiogenesis (Algenstaedt et al., 2003, Galiano et al., 2004, Nieuwdorp et al., 2006), and (5) impaired metabolism of extracellular matrix molecules, such as hyaluronan (Huang and Kyriakides, 2020, Shakya et al., 2015). In addition, hyperglycemia leads to generation of advanced glycation end products (AGEs), vasculopathy, and hypoxia, which cause toxicity to keratinocytes and fibroblasts and exacerbate inflammation in the wound, thereby hindering regeneration of a proper extracellular matrix (ECM) (Dinh et al., 2012, Fang and Lan, 2023, Shaikh-Kader et al., 2019). In a study examining diabetic patients with chronic foot ulceration, patients who failed to heal had higher serum levels of proinflammatory cytokines and more inflammatory cells in their wound biopsies, compared to patients who ultimately healed (Dinh et al., 2012). Many other reports also support the existence of a persistent, pro-inflammatory state in diabetes (Donath and Shoelson, 2011).

Tumor necrosis factor stimulated gene 6 (TSG-6) was first identified as a gene induced by TNFα (Lee et al., 1990). TSG-6 protein is secreted by many cells in response to injury, and binds to a wide range of ligands including glycosaminoglycans (GAGs) and the extracellular matrix proteins aggrecan, fibronectin (FN), and thrombospondin-1 (TSP1). TSG-6 also interacts with serum inter-alpha-inhibitor (IαI) and pre-alpha-inhibitor (PαI), and several growth factors (Day and Milner, 2019, Milner and Day, 2003). Through its link module, TSG-6 can bind to cytokines and chemokines (e.g., CXCL8, CXCL11, and CCL5), preventing them from interacting with GAGs on the endothelial surfaces of small blood vessels; thus TSG-6 can hinder proper chemokine presentation to leukocytes and prevent chemotaxis and trans-endothelial migration (Day and Milner, 2019, Dyer et al., 2014). Conversely, a reduced level of TSG-6 in the skin should allow more free cytokines to bind to endothelial-associated GAGs, enhance leukocyte chemotaxis, and significantly alter the overall landscape of leukocyte-generated cytokines at the wound site.

In an earlier study on the role of TSG-6 in wound healing, we showed that TSG-6 gene deletion leads to delayed epithelial closure, elevated TNFα expression, abnormal persistence of neutrophils, and delayed resolution of the granulation phase of wound healing in TSG-6 knockout mice (Shakya et al., 2020). Those findings established that a loss of TSG-6 expression can adversely affect inflammatory and matrix regenerative processes required for normal wound healing. Here, we report that TSG-6 expression is substantially reduced in diabetic skin wounds, suggesting that low TSG-6 levels in diabetic skin may be functionally linked to delayed wound healing in diabetes. We investigated this potential association in monogenic *db/db* mice and also in a diet- and obesity-associated murine model that more closely resembles human T2DM. In this report we show that TSG-6 expression is significantly reduced in both wounded and unwounded diabetic skin, and that abnormal closure and changes in cytokine expression are reversible by re-introduction of TSG-6 into the wound.

## RESULTS

### Genetically obese diabetic mice (*db/db*) show delayed wound closure and reduced TSG-6 expression

As a first approach, we studied wound healing in Lepr^db/db^ (“*db/db”)* mice, a well-established model of severe obesity and hyperglycemia due to a mutation in the Leptin receptor and featuring delayed cutaneous wound closure (Huynh et al., 2020). To assess TSG-6 expression after injury, we created 5 mm punch-biopsy wounds in *db/db* and wildtype control mice as described (Wang et al., 2013); wounds were secured with silicone splints to reduce variability caused by dermal contracture (**Figure 1a, b**). As expected, wound closure was significantly delayed (**Figure 1c)**. In parallel experiments, TSG-6 levels in unwounded skin were compared to those in wounded skin at Day 1 or Day 12 post-wounding (for both *db/db* mice and wildtype mice) using Western analysis (**Figure 1d)**. After wounding, TSG-6 expression was induced in both wildtype mice and *db/db* mice, but the magnitude of induction was relatively less in *db/db* wounds compared to wildtype controls (**Figure 1e)**.

**Figure 1.**
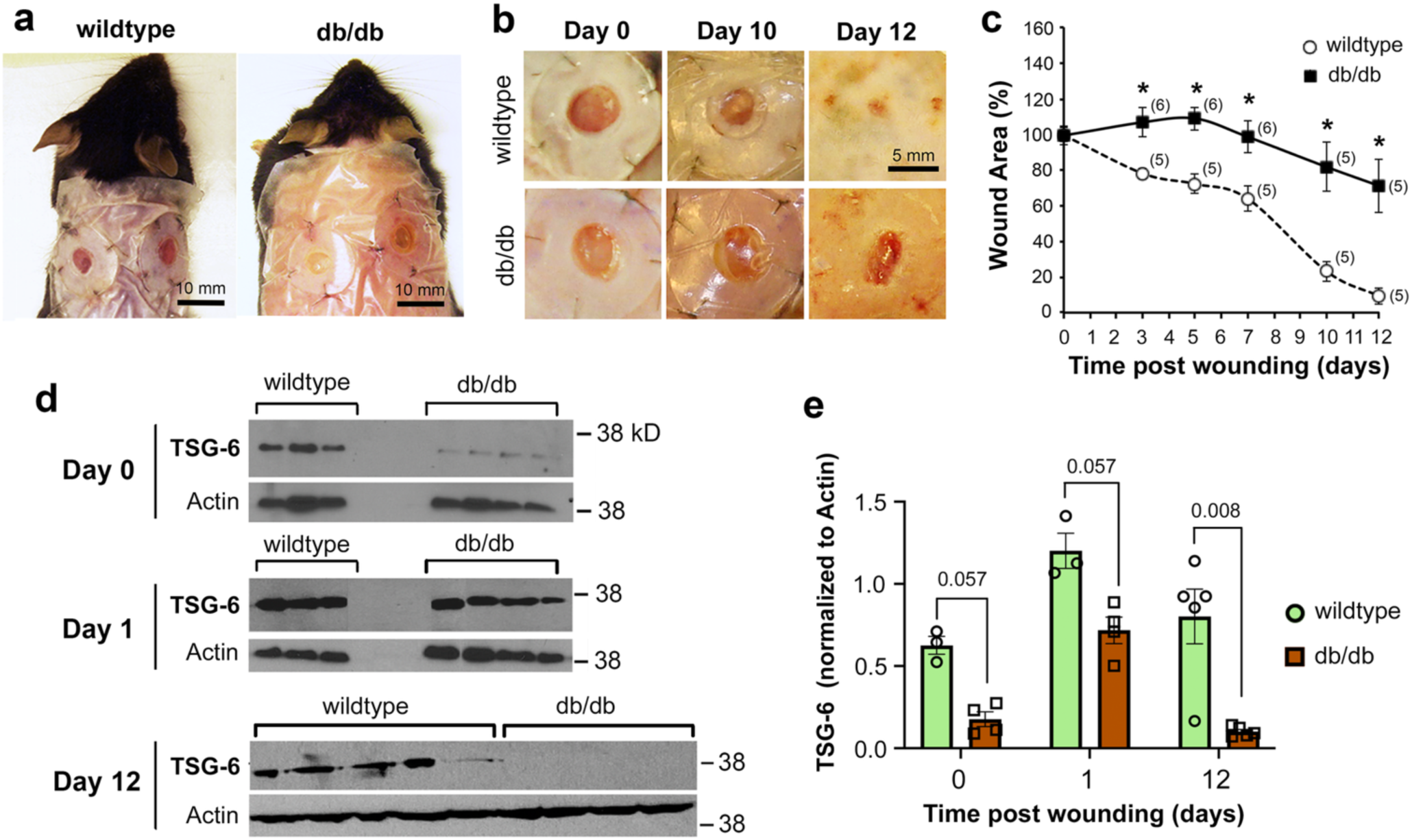
Wound closure is delayed and expression of TSG-6 is reduced in *db/db* mutant mice. For these experiments, mice were fully anesthetized, the fur removed by shaving and depilation, and full-thickness excisional wounds created with a 5-mm biopsy punch; the wound site was then secured with silicone splints. (**a**) *Lepr ^db/db^* mice (*db/db*) appeared grossly obese relative to the controls (C57BLKS/J strain; *wildtype*). (**b**) Photographic examples of wounds on days 0, 10, and 12. (**c**) Quantification of wound area over a 12-day time course. *Parentheses,* number of mice per time point. *Asterisks*, difference between db/db and control was significantly different, per Student t-test, p < 0.001. *Note:* On day 0, blood glucose levels were 185 ± 35 mg/dL (mean ± SD) for control mice, and >600 mg/dL for db/db mice (above the glucometer limit of detection). (**d**), Western blot analysis of wound skin at 0, 1, and 12 days post-injury in db/db mice or C57BLKS/J controls (*wildtype*); 3 to 5 mice per time point are shown. (**e**) Quantification of the TSG-6 protein bands from Westerns; *numbers above the bars* are P values from Mann-Whitney nonparametric test.

A problem with the *db/db* genetic model is that fasting blood glucose levels are very high (e.g., >600 mg/dL in our 8-week-old mice), vastly exceeding the levels of sustained hyperglycemia typically observed in human T2DM diabetic patients (Nichols et al., 2008). This was inconsistent with our goal of investigating TSG-6 in a model that more closely resembles human T2DM physiology (which features a multifactorial rather than monogenic etiology, with relatively moderate levels of hyperglycemia).

### Development of a murine model of T2DM with consistent hyperglycemia and ability to survive the stress of wounding

To induce T2DM, we first tried a diet-only approach. C57BL/6J mice were maintained on a high-fat diet for an extended period of 8 months (**Supp Table S1; Supp Fig. S1**). Although mice on the high fat diet weighed ∼40% more than control mice throughout that period (**Supp Fig. S1a**), they failed to become hyperglycemic, never reaching the diabetic threshold of 250 mg/dL (**Supp Fig. S1b**). When splinted wounding was performed with these mice, no significant difference in closure rates between the groups was seen (**Supp Fig. 1c, d**).

A literature search for alternative models of diabetes in mice (**Supp Table S2**) produced a body of evidence suggesting that low-dose streptozotocin (STZ), a glucose analog that kills pancreatic beta cells, could be a useful approach in combination with a high fat diet. However, wide variability hindered applicability of the prior reports. For example, most reports dealt with Type 1 diabetic wound models wherein mice receive STZ injections while on a normal diet (An et al., 2020, Li S. et al., 2022, Wang et al., 2019). However, for T2DM there was only one study that described skin wounding in C57BL/6 mice exposed to a high-fat diet plus STZ (Sun et al., 2020). In that report, mice received one injection of 40 mg/kg STZ and failed to develop a significant elevation in blood glucose levels (Sun et al., 2020). After considering all prior studies that used T2DM models (**Supp Table S2**) we performed pilot experiments using a range of STZ dosing regimens, starting with a single 100 mg/ml dose of STZ and progressing to lower dose protocols. The higher doses proved excessively toxic; mice experienced substantial weight loss and became too sick to withstand subsequent wounding. Lower doses (two daily injections of 40 mg/mL) failed to consistently induce hyperglycemia within the experimental time window. Ultimately, the dosing strategy was adjusted to deliver three consecutive days of STZ at 40 mg/kg, leading to our final protocol (**Fig. 2a**). With this optimized regimen, mice achieved significant obesity and did not lose weight post-STZ (**Fig. 2b**). By one-week post-injection, consistent hyperglycemia was evident from a high fasting blood glucose level (**Fig. 2c**) and an abnormal glucose tolerance test (**Fig. 2d, e**). In addition, an insulin tolerance test (ITT) also indicated impaired glucose homeostasis (**Supp Fig. S2**). Although blood glucose levels decreased initially after insulin injection in the HFD+STZ mice, blood glucose levels later remained elevated (the same as the HFD-only group), indicating insulin resistance.

**Figure 2.**
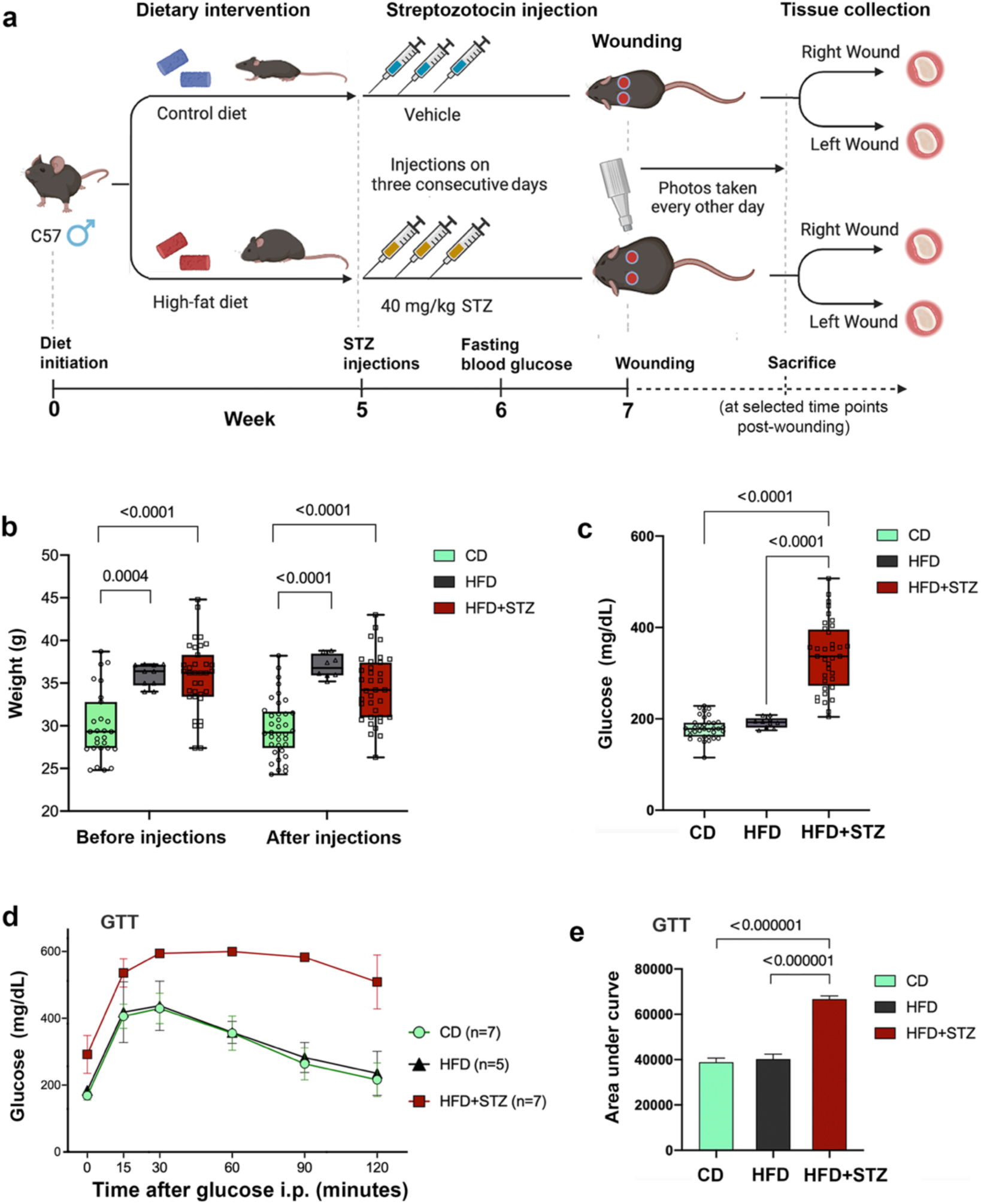
Creation and characterization of an obese diabetic wound model that combines a high-fat diet (HFD) with streptozotocin (STZ) injections. (**a**) Schematic of protocol to induce diabetes. (**b**) Body weights of the mice before and at one week after STZ injections. P-values are shown above the bars for each comparison from the non-parametric Mann-Whitney test. (**c**) Fasting blood glucose levels one week after completion of STZ injections. (**d**) Glucose tolerance test (GTT), showing the time course of blood glucose levels after a glucose bolus at time zero; (**e**) Area under the Curve for the GTT results; P-values are from student t-test; results of the one-way ANOVA revealed significant differences among these three groups, whereas Post-hoc Tukey’s HSD tests indicated significant differences between CD and HFD+STZ and between HFD and HFD+STZ, but not between CD and HFD (p = 0.85).

### Wound closure is delayed and TSG-6 expression is reduced in HFD/STZ-induced diabetic mice

To effectively capture the impact of elevated glucose levels and avoid possible recovery of β-cell function, wound healing experiments were performed within 10-14 days after STZ injection. These mice remained relatively healthy (**Fig. 3a**) with normal survival rates after wounding. Notably, HFD+STZ mice showed impaired wound closure (**Fig. 3b**). At Day 10, for example, the wound area (% of initial size) in control mice was 16.5 ± 2.3, *versus* 35.8 ± 3.7 in diabetic mice (mean ± SEM). The full-time course study showed that diabetic closure rates were statistically different (delayed) at 5, 7, and 10 days post-wounding (**Fig. 3c**).

**Figure 3.**
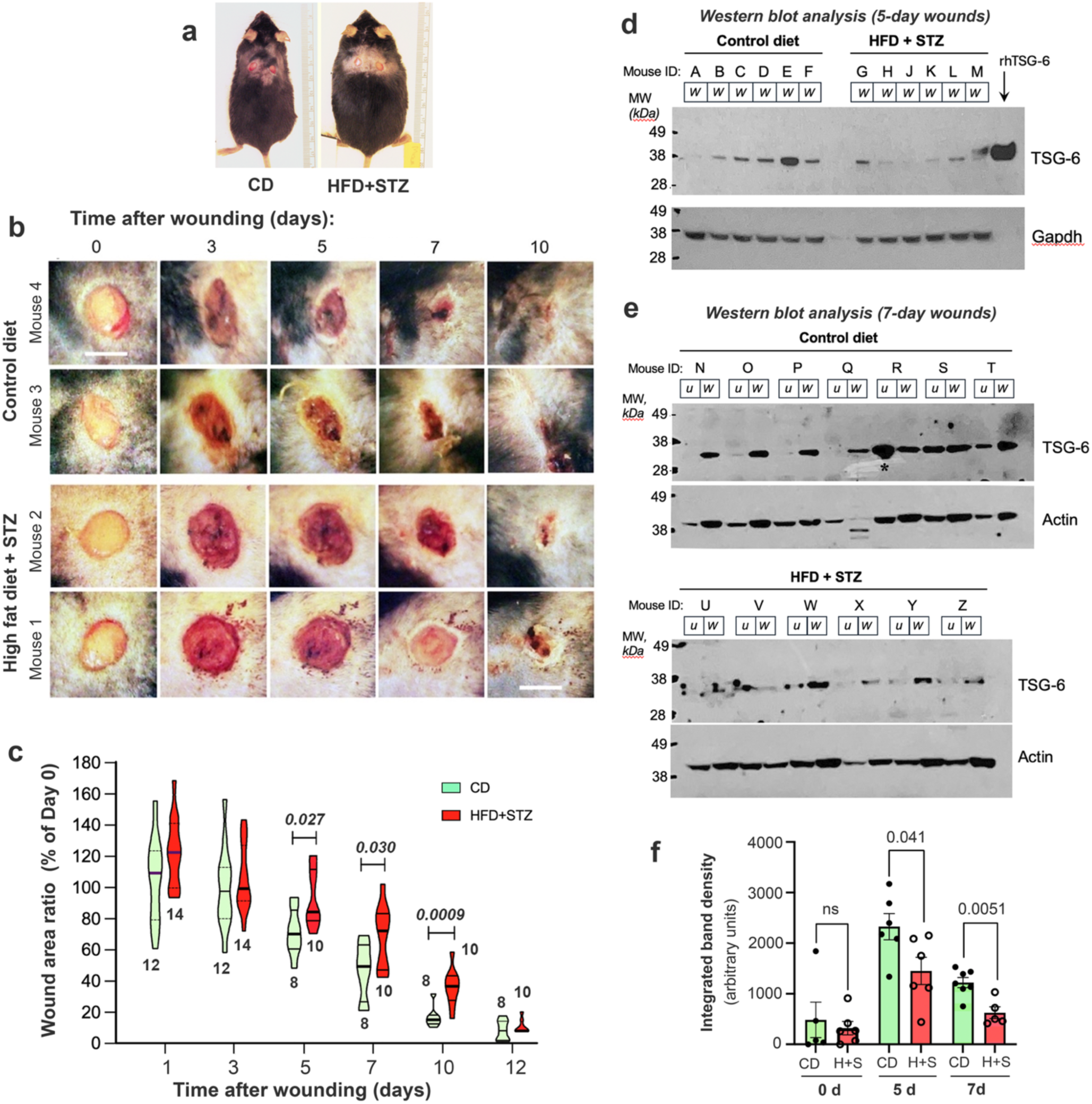
Decreased TSG-6 expression is associated with delayed wound closure in obese, diabetic HFD+STZ mice. (**a)** Photos comparing a typical control diet-fed mouse versus a HFD+STZ treated mouse, immediately after wounding under full anesthesia. (**b)** Examples of wounds from 4 different mice to illustrate the time course of wound closure. *Scale bar*, 5 mm. (**c**) Quantification of wound area, demonstrating delayed closure in HFD+STZ mice. Numbers of mice (beneath each symbol) and P-value of Mann-Whitney nonparametric test (above each symbol) are indicated. (**d**) TSG-6 protein expression in 5 day wounds. (**e**) TSG-6 protein expression in 7 day wounds (*W*), and in unwounded skin (*u*) paired from the same mouse; a total of 25 biopsied mice are shown. (**f**) Quantification of Westerns from multiple Western blots, normalized to loading controls. *CD,* control diet; *H+S,* high fat diet+STZ treated. P values from Mann-Whitney nonparametric tests are shown above brackets.

TSG-6 expression in wounds from these experiments was significantly less in the HFD+STZ diabetic mice at five days (**Fig. 3d**) and seven days (**Fig. 3e**) post-wounding (**Fig. 3f**). In addition, TSG-6 protein measurements by a second method, ELISA, confirmed that TSG-6 levels are lower in unwounded diabetic skin relative to intact skin of nondiabetic mice (**Supp Fig. S3**).

### Delayed wound closure in HFD/STZ diabetic mice is partially reversed by re-introduction of TSG6

In earlier work, we had shown that delayed wound healing in TSG-6 knockout mice could be reversed by injecting recombinant TSG-6 protein into the wounds at Day 0 and Day 4 post-injury (Shakya et al., 2020). We used human recombinant TSG-6 (rhTSG-6) because mouse and human TSG-6 share 93-94% amino acid sequence homology, providing a strong functional similarity as shown in multiple preclinical studies wherein human TSG-6 bound successfully to murine extracellular receptors and functioned seamlessly in mouse models across several disease categories (Li R. et al., 2022). However, in our HFD+STZ diabetic mice, a two-dose regimen of rhTSG-6 failed to normalize the delay in healing compared to vehicle-only injections (**Supp Fig. S4**), possibly due to compensatory mechanisms not present in the knockout model. Therefore, we tried administering three injections (at Days 0, 2, and 4; see **Fig 4a**). In these new experiments, a significant rescue in wound closure rates was evident by Day 4 and Day 7, observed photographically (**Fig 4b**) and histologically (**Fig 4c**). Summary results for 7 mice (14 wounds per condition) are displayed graphically in **Fig 4d**, confirming a significant reversal of the wound closure delay.

**Figure 4.**
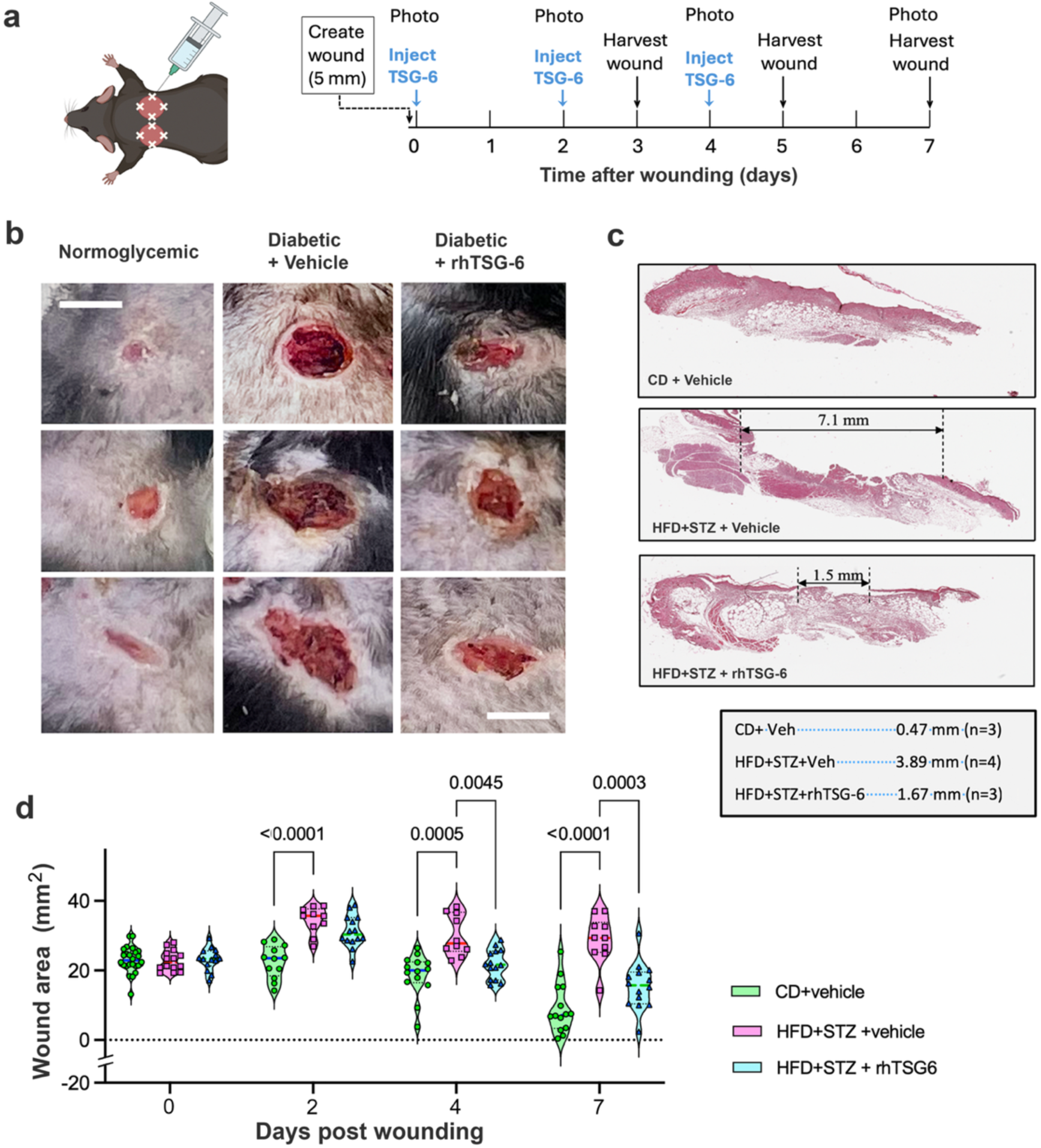
Delayed wound closure in HFD+STZ treated mice is partially reversed after injection of recombinant human TSG-6 (rhTSG-6). (**a**) Protocol used for intralesional injections of rhTSG-6. (**b**) Examples of wounds from 9 different mice, all photographed at 7 days post-injury. *Scale bar,* 5 mm. (**c**) Histologic examination of wounds harvested at 7 days post-injury; length of the intraepithelial wound gap is illustrated in H&E stained images, and mean values are shown in the table. (**d**) Time course of wound closure; photographs were taken at days 0, 2, 4, and 7 post-injury and open wound area determined using NIH ImageJ software. n = 14 wounds/condition; P values for Mann-Whitney test are shown above the bars.

### Diabetes-related differences in inflammation-regulated cytokine expression are reversed by re-introduction of TSG-6 into diabetic wounds

To support our hypothesis that the low TSG-6 levels observed in diabetic skin contribute to abnormal wound healing through loss of TSG-6 associated anti-inflammatory functions, we surveyed diabetic wound tissues for changes in cytokine expression (**Fig. 5**). Cytokines with known pro-inflammatory properties (IL-6, KC, MIP-1α, and MCP-1) were increased in HFD+STZ treated mice after wounding to a greater extent than in normoglycemic control mice (**Fig 5a, c, d, e**). Notably, the magnitude of these inductions was less when rhTSG-6 was added to the regimen (observed at Day 7 for IL-6, KC, and MIP-1α, and also at Day 5 for IL-6). The anti-inflammatory cytokine IL-10 showed opposite behavior, i.e., wounding-induced IL-10 levels were suppressed in HFD+STZ mice at Days 5 and 7 but restored by the re-introduction of rhTSG-6 (**Fig 5b**). For two other cytokines, VEGF and TNFα, only slightly increased levels were seen at Days 3 and 5 post-wounding, but by Day 7 there was a much larger induction in the HFD+STZ mice relative to normoglycemic mice, and this was reversible by injections of rhTSG-6 (**Fig 5f, g**). We conclude that wounding in HFD+STZ diabetic mice generates higher levels of pro-inflammatory cytokines and lower levels of an anti-inflammatory cytokine (IL-10) as compared to non-diabetic wounds; re-introduction of rhTSG-6 reverses many of those abnormalities.

**Figure 5.**
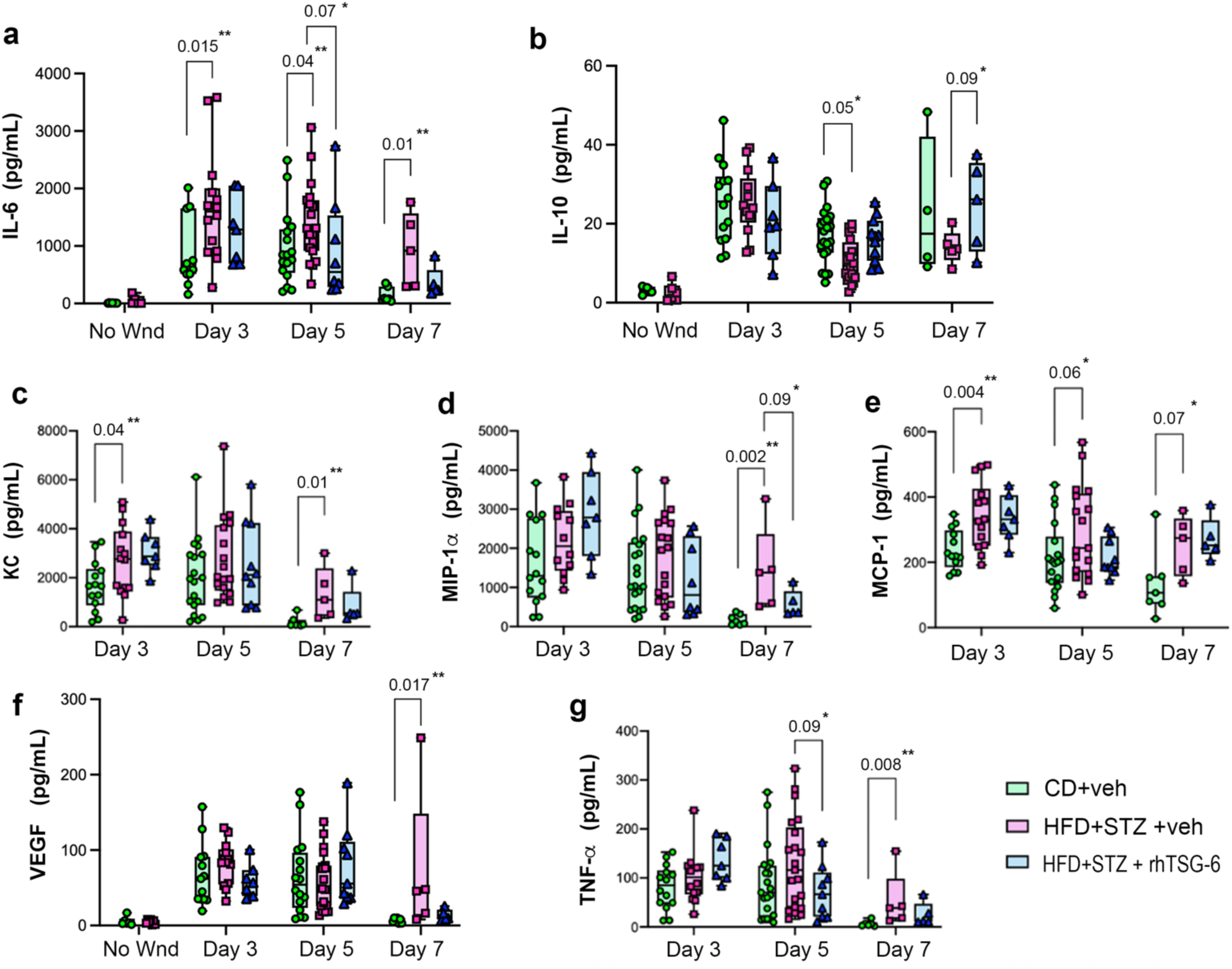
Cytokine and chemokine profiles in control and HFD+STZ mice at various times after wounding, with or without administration of rhTSG-6. Panels show the levels of (**a**) IL-6, (**b**) IL-10, (**c**) KC, (**d**) MIP-1α, (**e**) MCP-1, (**f**) VEGF, and (**g**) TNF-α, as measured by multiplex bead immunofluorescent assay before and after wounding. Statistical analyses included the Kolmogorov-Smirnov test for normality, followed by either a Student’s t-test or the Mann-Whitney test. Consistent with the exploratory nature of this study, P-values above the bars indicate changes that are significant at the (*) P< 0.1 or (**) P < 0.05 level.

### Profiling of immunocytes in wounds reveals diabetes-related effects upon macrophage recruitment

To investigate whether Type 2 diabetes and TSG-6 might affect the orchestrated recruitment of inflammatory cells into murine wounds, an experiment using the protocol in **Fig. 4a** with biopsy of wound tissues at Days 3 and 7 was conducted. Paraffin-fixed tissues were sectioned and analyzed via immunohistochemistry for the presence of inflammatory cells using antibodies specific for neutrophils, macrophages, and T cells (see **Supp Methods**). Employing image software-assisted techniques to identify and count each leukocyte cell type within the wounds, no relative difference in neutrophils or T-cells was observed (**Supp Fig. S5**). However, macrophages displayed interesting differences. At Day 7, diabetic (HFD+STZ) wounds had more F4/80-stained dermal macrophages than in normal mouse wounds, and injection of rhTSG-6 increased these macrophage numbers (**Fig. 6a**). Macrophages in subcutaneous fat, which tend to cluster around adipocytes in a characteristic arrangement called “crown-like structures (CLS)” (Chini et al., 2026, Murano et al., 2008), showed higher numbers of CLS in HFD+STZ mice, which was further augmented when rhTSG-6 was injected (**Fig. 6b**). These findings suggest that diabetes causes dysregulation of overall macrophage recruitment in both the dermal and subcutaneous wound compartments, and that TSG-6 re-introduction may selectively enhance recruitment of one or more subsets of macrophages in the dermis and subcutaneous fat.

**Figure 6.**
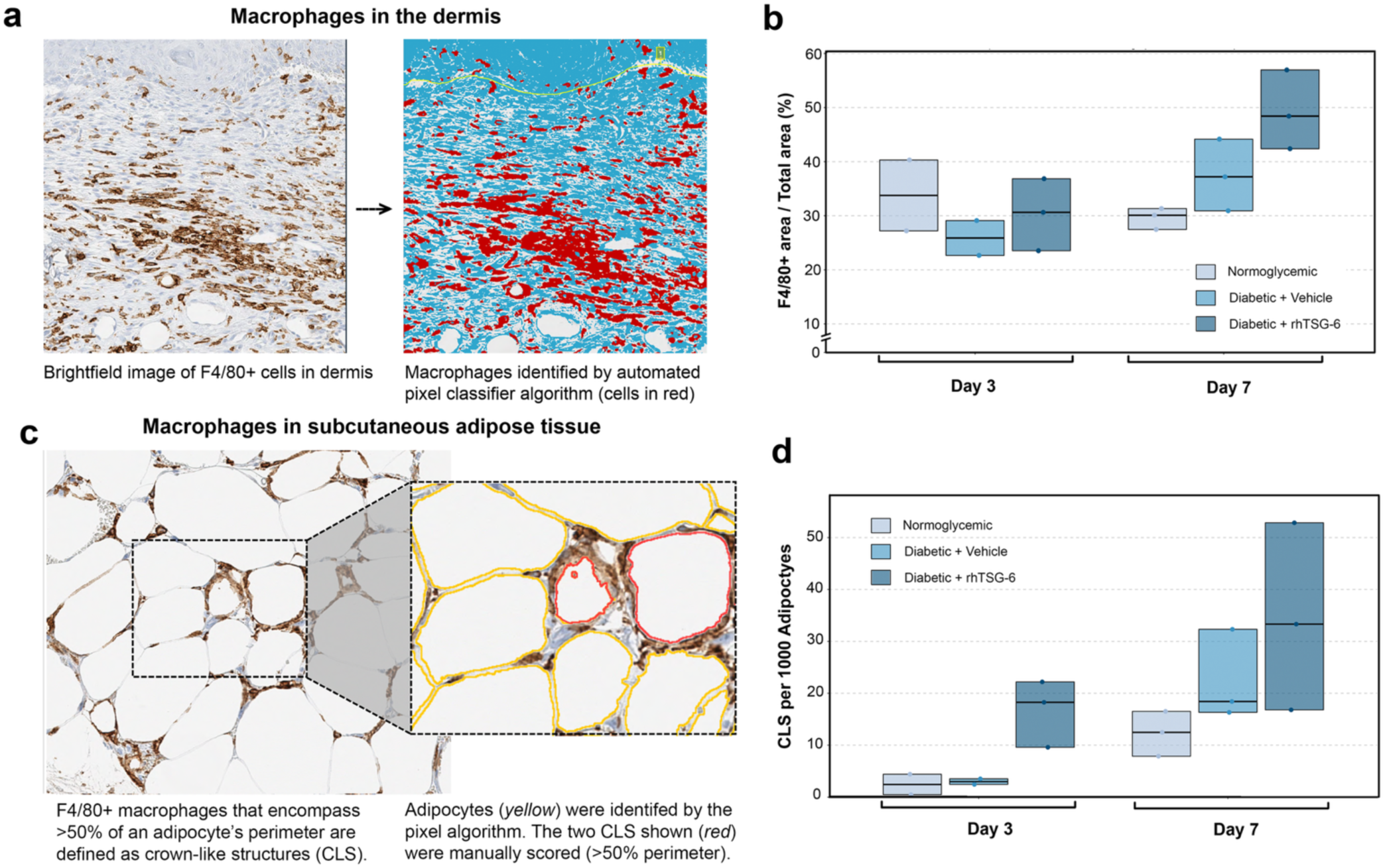
Macrophage recruitment in 7-day wounds is elevated in diabetic HFD+STZ mice, and further increased by injection of rhTSG-6. Histological sections of 3-day and 7-day old wounds were immunostained with F4/80 antisera to label macrophages, which were then analyzed using Q-path software as described in Supplemental Methods. (**a, b**) *Analysis of macrophages (*in the dermis beneath the wound bed; (**a)** Illustration of semi-automated cell count analysis; (**b**) Graphical summary of dermal macrophage counts under normal and diabetic conditions. (**c, d**) *Analysis of crown-like structures (CLS)* in the subcutaneous fat beneath wounds; (**c**) Image example showing how CLS were scored; (**d**) Graphical summary of CLS counts normalized to adipocyte number in wounded control and diabetic skin.

## DISCUSSION

In this study, we investigated the hypothesis that TSG-6 exerts an overall suppressive effect upon certain aspects of inflammation in wounded diabetic skin, where TSG-6 protein is found to be relatively lacking. In two different mouse models (monogenic *db/db* mice, and high fat diet/streptozotocin mice), we showed that TSG-6 expression is significantly lower in diabetic skin than in normoglycemic skin, both prior to wounding and after wounding. Wound closure rates were significantly delayed in both T2DM models. The notion that low TSG-6 levels in diabetic skin may be partially responsible for delayed wound closure was supported by our demonstration that reintroduction of TSG-6 protein into diabetic wounds partially reversed the delay in wound healing. Furthermore, abnormal changes in cytokine expression were observed in T2DM wounds (over-induction of proinflammatory IL-6, KC, MIP-1alpha, VEGF, and TNFα; repression of anti-inflammatory cytokine IL-10). The fact that loss of TSG-6 disrupts cytokine levels within healing wounds, yet those abnormalities are reversed when TSG-6 protein levels are restored, indicates that TSG-6 may represent an important regulator of the inflammatory cytokine landscape during wound healing.

An important goal in this study was to try to emulate human T2DM wound healing, a multifactorial process that cannot be realistically portrayed in a single-gene mutant model (the *db/db* mouse). Because T2DM is obesity-related, we initially attempted to replicate an experimental system in C57BL/6J mice (Surwit et al., 1988) in which obesity, hyperglycemia, and elevated insulin levels were achieved (**Supp Table S2**, first entry). Using that same model, Seitz (Seitz et al., 2010) had reported delayed wound closure in obese, HFD-fed mice (**Supp Table S2**, second entry), but when we attempted to do the same, glucose levels did not reach diabetic thresholds and delayed wound closure was not observed, perhaps because our mice were not hyperglycemic. Others have reported similar negative results and lack of hyperglycemia in HFD-only C57BL/6J mice (Tierney et al., 2022).

Subsequently, we searched the literature to find an alternative, practical Type 2 diabetic model (**Supp Table S2**). A point generally not addressed in studies with diabetic mice (most of which involve Type 1 diabetes induced by STZ injection) is the fact that mice on a normal diet typically lose weight after STZ, become very sick, and cannot survive the stress of wounding. While this problem is less common with Type 2 diabetic models, only a handful of published reports examined cutaneous wound healing. Experimental conditions in those reports varied so widely that it became necessary for us to test several of them. Ultimately, we chose the regimen described by Gilbert et al. (Gilbert et al., 2011) in which HFD-fed mice receive relatively low nontoxic doses of STZ over three days, allowing them to develop diabetes yet survive the wounding stress. Notably, we found we could eliminate silicone splints from the protocol and still measure reproducible effects upon wound closure. This was very helpful because mice tend to gnaw and destroy their splints, creating additional skin inflammation that confounds interpretation of inflammatory cytokine changes in wounds.

Mechanistically, we had previously established a strong association between TSG-6 levels and wound closure in TSG-6 null mice (Shakya et al., 2020). In the current report, we have extended the association between TSG-6 and wound healing to a clinically relevant situation (T2DM) in which diabetic mice display a partial loss of TSG-6 protein in their skin. We showed that the reduction in TSG-6 is functionally linked to a pro-inflammatory state within the diabetic wound. Our survey of wounding-related cytokines, while not exhaustive, showed normalization of expression of 6 out of 7 cytokines following TSG-6 re-introduction (**Fig. 5**), providing strong evidence for an anti-inflammatory effect of TSG-6.

To explore possible cellular sources for these TSG-6 regulated cytokines, we interrogated the inflammatory cell landscape in wounds using immunohistochemical staining. Our survey identified a diabetes-related increase in overall F4/80+ macrophages within the dermis and subcutaneous adipose tissue of HFD+STZ wounds at 7 days, relative to control wounds (**Fig. 6**). Interestingly, while more macrophages were present in the dermis and subcutaneous fat of diabetic (HFD+STZ) mice, re-introduction of rhTSG-6 into HFD+STZ wounds caused an even further increase in overall macrophage numbers. The latter observation is difficult to reconcile with reduction in pro-inflammatory cytokine levels observed in **Fig. 5**, unless the macrophages recruited into the diabetic + rhTSG-6 wounds have a tissue-repair phenotype (M2) which secrete less pro-inflammatory cytokines than M1 macrophages. In fact, such a scenario was reported in a study on corneal epithelial wound healing in diabetic mice, wherein injections of TSG-6 caused a “macrophage switch” that correlated with reduced inflammation and accelerated corneal wound closure (Di et al., 2017). If future resources allow, we hope to be able to conduct experiments to conclusively determine whether a similar macrophage phenotype switch occurs in diabetic skin wounds before and after TSG-6 restoration.

## MATERIALS AND METHODS

Full details about the methods used in this paper are available in **Supplementary Materials**.

### Animals

C57BLKS/J and Lepr^db/db^ (BKS.Cg-Dock7+/+Lepr db/J) male mice, and C57BL/6J mice for the dietary+STZ experiments, were purchased from Jackson Laboratories (JAX; Bar Harbor, ME).

### Diets and treatments

The two diet formulations employed here (high-fat diet, Cat. #D12331; control diet, Cat. # D12329; from Research Diets, Inc., New Brunswick NJ) are listed in **Supp Table S1**. Mice in the HFD+STZ group received STZ (40 mg/kg i.p.) daily for 3 days. Tail vein blood was obtained for measurements of fasting blood glucose, GTT, and ITT at 7 days after initial STZ injection. Wounding studies were performed ∼10 days after initial STZ injection. All procedures were approved by the Cleveland Clinic Institutional Animal Care and Use Committee (IACUC).

### Glucose tolerance test (GTT)

After fasting for 6 hours, basal glucose levels were measured in tail vein blood using a standard glucometer. A glucose solution (1.5 g per kg) was injected i.p., and blood glucose measured at 15, 30, 60, 90, and 120 minutes.

### Insulin resistance test (ITT)

After a 6-hour fast, basal glucose levels were measured. Insulin (0.75 IU per kg body weight) was injected i.p., and glucose levels measured at 15, 30, 60, 90, and 120 minutes. Wounding (Non-splinted protocol): C57BL/6J mice were shaved 24-48 hours prior to wounding. On the day of wounding, a 5-mm biopsy punch was used to create two full-thickness wounds. Closure was monitored using a digital camera; wound areas were determined using NIH Image J software.

### Wounding (Splinted protocol)

For *db/db* mice and matched-strain wildtype mice, wound splints were used as described (Wang et al., 2013).

### Recombinant TSG-6 injections

On days indicated in Fig. 4, recombinant human TSG-6 (rhTSG-6, R&D Systems, Minneapolis, MN) was injected by distributing a total of 2 μg in 100 μl PBS at five sites per wound. Control mice received PBS alone.

### Preparation of tissue lysates

After euthanasia, skin tissues were flash-frozen in liquid nitrogen, weighed, crushed in a tissue pulverizer on dry ice, homogenized in lysis buffer (1M Tris-HCl @ pH 7.5, 5M NaCl, 0.05%Tween-20, with protease inhibitors), and the supernatants stored at -80°C.

### Western blots

Lysates were separated on SDS-PAGE gels, blotted onto polyvinylidene membranes, and probed overnight at 4°C with one of the following primary rabbit antibodies: (1) TSG-6; (2) Actin; (3) GAPDH (see **Supp Methods** for sources). After washing, membranes were incubated with secondary goat anti-rabbit HRP and developed using a chemiluminescent ECL reagent. Blots were stripped and re-probed for actin or GAPDH as loading controls.

### Enzyme-linked immunosorbent assay (ELISA)

TSG-6 protein levels were quantified using the TSG-6 ELISA Kit (Mouse TNFAIP6 / TSG-6 ELISA Kit from LSBio) per manufacturer’s instructions.

### Multiplex cytokine assay

Cytokine levels were quantified using the Mouse Cytokine Magnetic Kit (Catalog ID: MCYTOMAG-70K-07 from Millipore-Sigma), per manufacturer’s instructions.

### Immunohistochemistry and digital analyses of immune cell populations

Skin tissues were fixed in Histochoice, paraffin-embedded, cut into 5-micron sections, and immunostained for neutrophils, macrophages, and T-cells. Inflammatory cell populations in wounds were enumerated using QuPath software algorithms as described in **Supp Materials.**

### Statistical analysis

Data were analyzed using GraphPad Prism 9.0.0 software. Results are presented as means ± SEM. If data passed tests of normality, then Student’s t-test for statistical significance was used; otherwise, Mann-Whitney tests were employed. The significance threshold was set at P < 0.05.

