## Supplemental Tables, Figs, & Methods for "Abnormally low expression of tumor necrosis factor stimulated gene 6 (TSG-6) in obese type 2 diabetic mice contributes to altered inflammation and delayed cutaneous wound healing"

### Supplementary Table S1. Formulations of the High Fat diet and Control diet.

#### (A) Formulation of the High Fat Diet

| PRODUCT # D12331 | INGREDIENTS | GRAMS% | KCAL% |
| --- | --- | --- | --- |
| **PROTEIN** | Casein, Lactic, 30 Mesh | 22.8 | 16.4 |
|  | Methionine, DL | 0.2 | 0 |
| **CARBOHYDRATE** | Sucrose, Fine Granulated | 17.5 | 12.6 |
|  | Maltodextrin 10 | 17 | 12.2 |
| **FAT** | Coconut Oil, Hydrogenated 101 | 33.4 | 54.0 |
|  | Soybean Oil, USP | 2.5 | 4.0 |
| **MINERAL** | Mix.S10001A | 4 | 0 |
|  | Sodium Bicarbonate | 1.05 | 0 |
|  | Potassium Citrate, Monohydrate | 0.4 | 0 |
| **VITAMIN** | Choline Bitartrate | 0.2 | 0 |
|  | V10001C | 1 | 0.7 |
| **DYE** | Dye, Red FD&C #40, Alum. Lake 35-42% | 0.01 | 0 |
| **TOTAL** |  | **100** | **100** |

#### (B) Formulation of the Control Diet

| PRODUCT # D12329 | INGREDIENTS | GRAMS % | KCAL% |
| --- | --- | --- | --- |
| **PROTEIN** | Casein, Lactic, 30 Mesh | 22.8 | 16.4 |
|  | Methionine, DL | 0.2 | 0 |
| **CARBOHYDRATE** | Sucrose, Fine Granulated | 47 | 60.1 |
|  | Maltodextrin 10 | 17 | 12.2 |
| **FAT** | Coconut Oil, Hydrogenated 101 | 4 | 6.5 |
|  | Soybean Oil, USP | 2.5 | 4.0 |
| **MINERAL** | Mix.S10001A | 4 | 0 |
|  | Sodium Bicarbonate | 1 | 0 |
|  | Potassium Citrate, Monohydrate | 0.4 | 0 |
| **VITAMIN** | Choline Bitartrate | 0.2 | 0 |
|  | V10001C | 1 | 0.7 |
| **DYE** | Dye, Red FD&C #40, Alum. Lake 35-42% | 0.01 | 0 |
| **TOTAL** |  | **100** | **100** |

##### Commercial Source:

Research Diets Inc., 20 Jules Lane, New Brunswick, NJ 08901, USA.

**Supplementary Table S2.** Representative studies along a timeline of development of HFD+STZ models to assess hyperglycemia and complications of Type 2 diabetes in C57BL/6J mice

| Author/Year | Doses of STZ;<br>(Interval<br>between<br>doses) | Mouse strain<br>(Vendor) | Age of<br>mice at<br>start of<br>HFD diet<br>(weeks) | Control diet<br>composition<br>(% wt/wt) | High fat diet<br>composition<br>(% wt/wt) | Duration on<br>HFD before<br>STZ was<br>given<br>(weeks) | Time after<br>STZ when<br>FBS was<br>measured<br>(weeks) | FBG<br>(mg/dL) on<br>control diet* | FBG<br>(mg/dL)<br>on high<br>fat diet<br>(HFD)** | FBG<br>(mg/dL)<br>after<br>HFD+STZ | T2DM<br>confirmed<br>by GTT<br>and ITT? | Delayed<br>wound<br>healing? |
| --- | --- | --- | --- | --- | --- | --- | --- | --- | --- | --- | --- | --- |
| Surwit, 1988 [1] | none given | C57BL/6J<br>(Jackson Labs) | 4 | fat 4.5%,<br>protein 23%,<br>carbo 56% | fat 36%,<br>protein 21%,<br>carbo 37% | **180 days,**<br>(HFD only) | (no STZ) | 169 ± 6.4 | 249 ± 8.2 | (no STZ) | Insulin<br>elevated | n/a |
| Seitz, 2010 [2] | none given | C57BL/6J<br>(Jackson Labs) | 6 | fat 11%,<br>protein 23%,<br>carbo 65% | fat 36.0%,<br>protein 23%,<br>carbo 36% | **190 days,**<br>(HFD only) | (no STZ) | 78 ± 5 | 90 ± 15 | (no STZ) | GTT (+);<br>insulin<br>elevated | YES |
| Luo, 1998 [3] | 1 dose of 100<br>mg/kg | C57BL/6J<br>(Jackson Labs) | 3 | fat 4.5%,<br>protein 23%,<br>carbo 46% | fat 36%,<br>protein 20%,<br>carbo 36% | 3 | 7 | 190 ± 5 | 253 ± 10 | 388 ± 38 | Insulin<br>elevated | n/a |
| Lian, 2007 [4] | 1 dose of 150<br>mg/kg | C57BL/6J<br>(Chinese Acad of<br>Sciences,<br>Shanghai) | 4 | fat 4.0%,<br>protein 21%,<br>carbo 47% | fat 26%,<br>protein 21%,<br>carbo 50% | 6 | 10 days | 105 ± 27<br>5.8 ± 1.5<br>mM | 159 ± 20<br>8.8 ± 1.1<br>mM | 366 ± 72<br>20.3 ± 4.0<br>mM | GTT (+);<br>ITT (+) | n/a |
| Gilbert, 2011 [5] | 3 doses of 40<br>mg/kg; (daily) | C57BL/6NCr (NCI<br>Frederick) | 24 | fat 4%,<br>protein 19%,<br>carbo 67% | fat 35%,<br>protein 26%,<br>carbo 26 % | 5 | 1 - 4 | 89 ± 19.7 | 104 ±<br>20.0 | 280 ± 10 | GTT (+);<br>ITT (+) | n/a |
| Mali, 2014 [6] | 3 doses of 40<br>mg/kg; (daily) | C57BL/6J (vendor<br>not stated) | 8 | Control diet<br>not specified | fat 35%,<br>protein 26%,<br>carbo 26 % | 2 | 4 | 265.2 ± 7.6 | nd | 540 ±<br>18.7 | GTT (+) | n/a |
| Yorek, 2015 [7] | 2 doses of 75<br>mg and 50 mg<br>(3 days apart) | C57BL6J<br>(Jackson Labs) | 13 | fat 4%,<br>protein 25%,<br>carbo 40% | fat 35%,<br>protein 26%,<br>carbo 26 % | 8 | 4 | 113 ± 8 | 163 ± 8 | 257 ± 18 | GTT (+) | n/a |
| Surbala, 2020 [8] | 4 doses of 35<br>mg/kg; (daily) | C57BL/6J (RIMS,<br>Manipur, India) | 7 - 9 | Standard<br>chow diet<br>(Pranav Agro<br>Industries Ltd,<br>Pune, India) | 58% of<br>calories from<br>fat | 10 | 1 | 120 ± 10 | nd | 335 ± 15 | GTT (+)<br>ITT (+) | n/a |
| Zhu, 2020 [9] | 5 doses of 40<br>mg/kg; (daily) | C57BL/6 (Tianqin<br>Biotech. Co.,<br>Hunan,<br>Changsha, China) | (20 ± 3<br>gm) | fat 5%,<br>protein 23%,<br>carbo 53% | fat 22%,<br>protein 20%,<br>carbo 48% | 8 | 1 | 75 ± 36<br>4 ± 2<br>mmol/L | nd | 414 ± 54<br>23 ± 3<br>mmol/L | GTT (+) | n/a |
| Sun, 2020 [10] | One dose of<br>40 mg/kg | C57BL/6 (Vital<br>River Lab Animal<br>Tech Co., Beijing,<br>China) | 5 | 10% of<br>calories from<br>fat | 60% of<br>calories from<br>fat | 16 | 2 | 124 ± 6 | nd | 210 ± 21 | GTT (+)<br>ITT (+) | YES |
| Alipour, 2026 [11] | 3 doses of 40<br>mg/kg (daily) | C57BL/6J<br>(Jackson Labs) | 8 | fat 7%,<br>protein 23%,<br>carbo 64% | fat 36%,<br>protein 23%,<br>carbo 34% | 5 | 1 | 180 ± 4.1 | 192 ± 3.6 | 337 ±<br>12.7 | GTT (+)<br>ITT (+) | YES *** |

### FOOTNOTES

\* FBG, fasting blood glucose, in mg/dL. Values that were originally reported as mmol/L (mM) are shown in italics. Conversion factor: 1 mg/dL = 0.0555 mmol/L.

\*\* FBG was measured just prior to STZ injection.

\*\*\* In this model of C57BL/6 mice, one can observe unequivocal wound closure delay with no splinting required; Clear development of hyperglycemia measurable by abnormal FBG, GTT and ITT, all in a total time of 7 weeks.

*carbo*, carbohydrates

*GTT*, glucose tolerance test

*HFD*, high fat diet

*ITT*, insulin tolerance test

*STZ*, streptozotocin

*n/a*, not assessed

### REFERENCES (cited in column 1 of the Supplementary Table)

1. Surwit, R.S., C.M. Kuhn, C. Cochrane, J.A. McCubbin, and M.N. Feinglos, "Diet-induced type II diabetes in C57BL/6J mice", *Diabetes*, 37(9) 1163-7. (1988).
2. Seitz, O., C. Schurmann, N. Hermes, E. Muller, J. Pfeilschifter, S. Frank, and I. Goren, "Wound healing in mice with high-fat diet- or ob gene-induced diabetes-obesity syndromes: a comparative study", *Exp Diabetes Res*, 2010 476969. (2010).
3. Luo, J., J. Quan, J. Tsai, C.K. Hobensack, C. Sullivan, R. Hector, and G.M. Reaven, "Nongenetic mouse models of non-insulin-dependent diabetes mellitus", *Metabolism*, 47(6) 663-8. (1998).
4. Lian, J., Y. Xiang, L. Guo, H. W., W. Ji, and B. Gong, "The use of High-Fat/Carbohydrate Diet-Fed and Streptozotocin-Treated Mice as a Suitable Animal Model of Type 2 Diabetes Mellitus", *Scan.J. Lab.Anim. Sci.*, 34(1) 21-29. (2007).
5. Gilbert, E.R., Z. Fu, and D. Liu, "Development of a nongenetic mouse model of type 2 diabetes", *Exp Diabetes Res*, 2011 416254. (2011).
6. Mali, V.R., R. Ning, J. Chen, X.P. Yang, J. Xu, and S.S. Palaniyandi, "Impairment of aldehyde dehydrogenase-2 by 4-hydroxy-2-nonenal adduct formation and cardiomyocyte hypertrophy in mice fed a high-fat diet and injected with low-dose streptozotocin", *Exp Biol Med* (Maywood), 239(5) 610-8. (2014).
7. Yorek, M.S., A. Obrosova, H. Shevalye, A. Holmes, M.M. Harper, R.H. Kardon, and M.A. Yorek, "Effect of diet-induced obesity or type 1 or type 2 diabetes on corneal nerves and peripheral neuropathy in C57Bl/6J mice", *J Peripher Nerv Syst*, 20(1) 24-31. (2015).
8. Surbala, L., C.B. Singh, R.V. Devi, and O.J. Singh, "Rutaecarpine exhibits anti-diabetic potential in high fat diet-multiple low dose streptozotocin induced type 2 diabetic mice and in vitro by modulating hepatic glucose homeostasis", *J Pharmacol Sci*, 143(4) 307-314. (2020).
9. Zhu, Y., C. Zhu, H. Yang, J. Deng, and D. Fan, "Protective effect of ginsenoside Rg5 against kidney injury via inhibition of NLRP3 inflammasome activation and the MAPK signaling pathway in high-fat diet/streptozotocin-induced diabetic mice", *Pharmacol Res*, 155 104746. (2020).
10. Sun, Y., L. Song, Y. Zhang, H. Wang, and X. Dong, "Adipose stem cells from type 2 diabetic mice exhibit therapeutic potential in wound healing", *Stem Cell Res Ther*, 11. (2020).
11. Alipour, M, et al. (This manuscript).

**Supplementary Figure S1.**

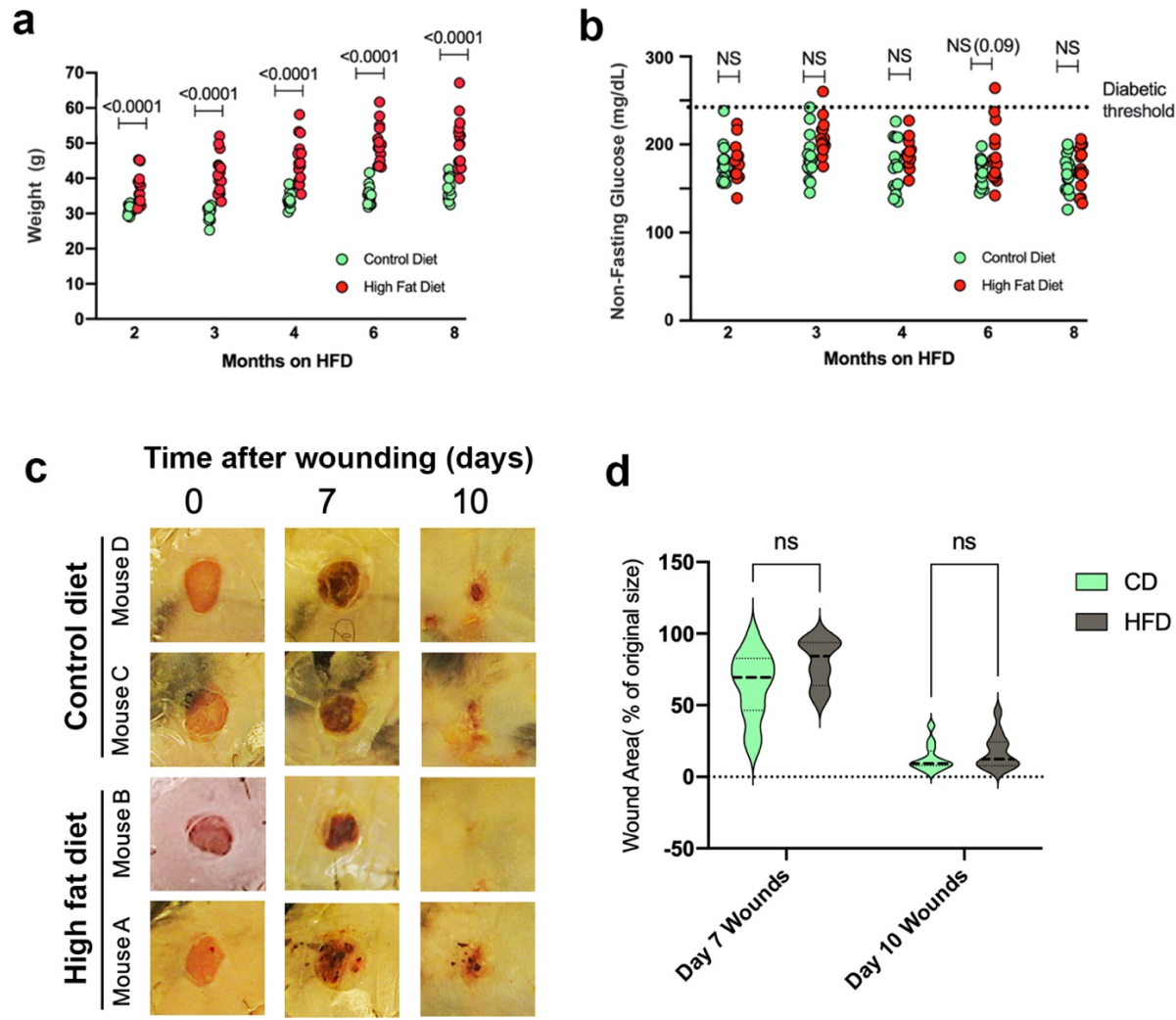

**Supplement Figure S1.** <u>Wound closure is not delayed in mice fed a high-fat diet with no additional intervention.</u> For eight months, C57/BL6 mice from JAX Laboratories were fed on a high-fat diet protocol (15 mice on HFD, 15 mice on CD). The mice became obese but did not develop hyperglycemia above the 250 mg/dL threshold. **(a)** Body weight (n=14 mice per group). Means ± SE; p-values are shown above the bars for each comparison from the non-parametric Mann-Whitney test. **(b)** Non-fasting glucose levels. **(c)** The C57/BL6 mice were subjected to the same wounding protocol as were the db/db mice in Figure 1. Representative images of four wounds at days 0, 7, and 10 post-wounding are shown. **(d)** Quantification of wound area for all mice at each condition, n = 12. (ns), no significant difference in wound size between the control mice and high-fat diet mice.

### Supplementary Figure S2.

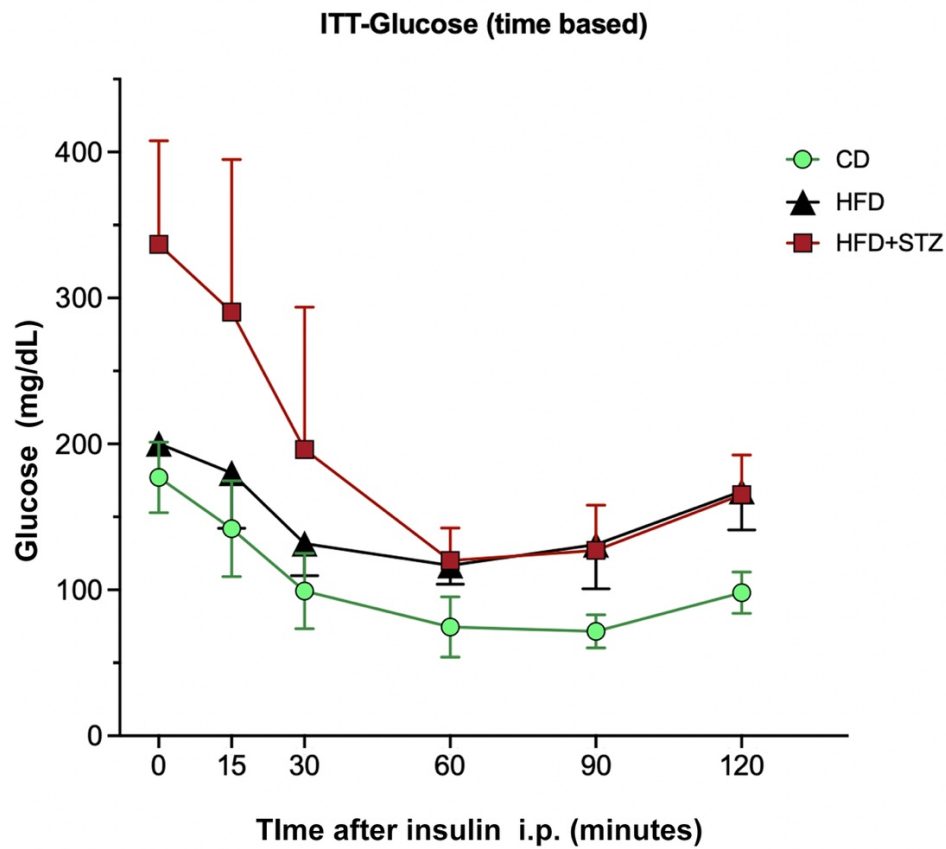

**Supplem Fig S2.** <u>Insulin tolerance test (ITT) reveals a biphasic response to glucose challenge in the HFD+STZ mouse model.</u>

**Technique:** Mice were placed in a clean cage with access to only water for a morning fast (6 hours; 8:00 am-2:00 pm). Mice were weighed, and an insulin solution was prepared for each mouse (0.75 IU insulin/kg body weight). If an animal became hypoglycemic with blood glucose below 20 mg/dL or appeared distressed, 300 μl of glucose solution (1.5 g/kg) was injected intraperitoneally (i.p.). Next, the basal level of glucose was measured using the glucometer. Then, insulin was injected via i.p., and glucose levels were measured in the tail vein blood after 15, 30, 60, 90, and 120 minutes.

**Result:** Statistical analysis of the data using a mixed-effects model with Type III fixed effects, revealed significant main effects of Time $F(2.041, 28.57) = 43.32$, $p < 0.0001$) and Diet $F(2, 15) = 13.96$, $p = 0.0004$), as well as a significant interaction effect (Time x Diet: $F(10, 70) = 5.490$, $p < 0.0001$). These omnibus ANOVA tests suggest overall differences across time points, diet groups, and their interaction. Post-hoc analyses using Tukey's test were performed to identify specific group differences indicated by p values.

**Interpretation:** The results of the ITT revealed an initial response to insulin in the HFD+STZ group, but blood glucose levels ultimately became resistant to insulin as they failed to drop to the same level as Controls, instead remaining similar to the HFD-only group (which has distinct insulin resistance). This indicates a comparable status of insulin resistance in the HFD-only and HFD+STZ groups.

#### Supplementary Figure S3

a

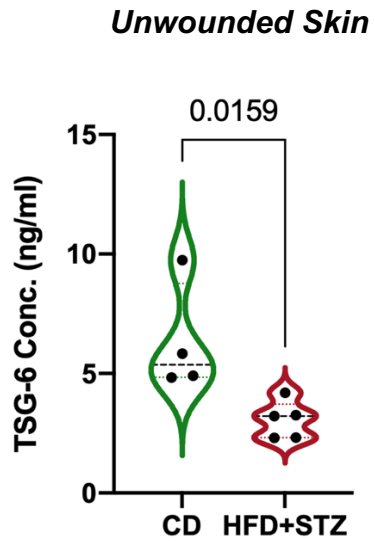

b

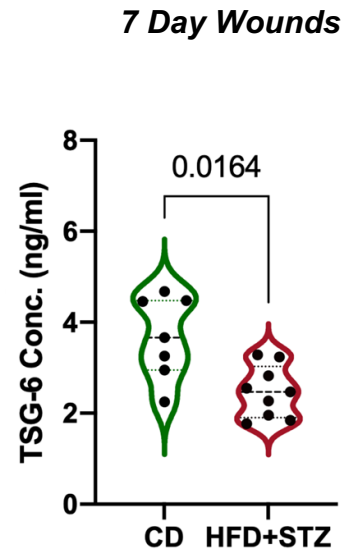

**Supplem Fig S3.** <u>TSG-6 concentration measured with ELISA assay.</u> Protein lysates derived from (a) Unwounded skin of C57 mice; (b) Wounded skin at 7 days post procedure. *Means* and *quartiles* are indicated by horizontal lines for each condition. *P-values* are from non-parametric Mann-Whitney test.

### Supplementary Figure S4

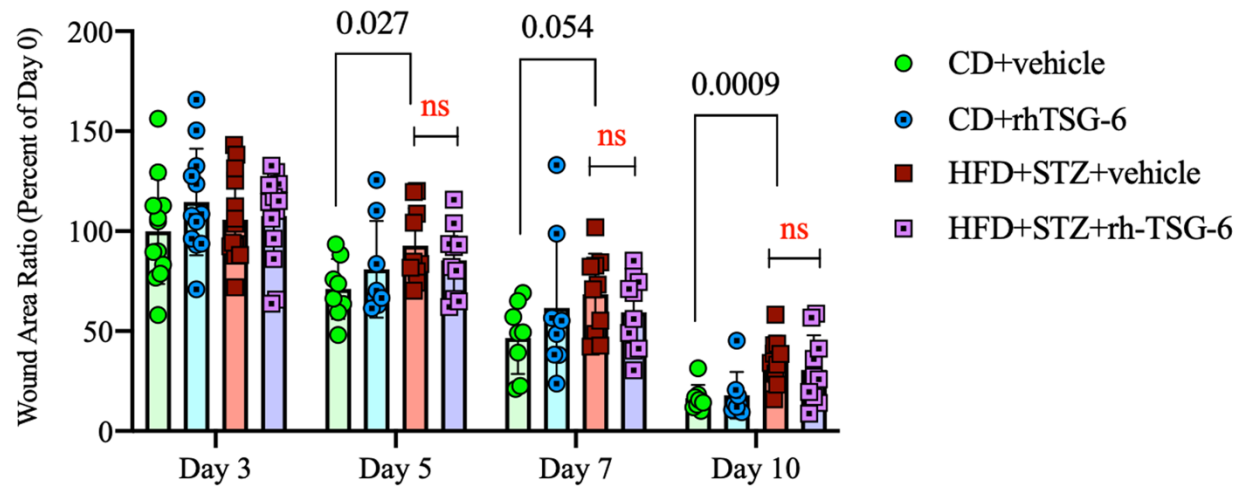

**Supplem Fig S4.** <u>A two-dose regimen of recombinant human TSG-6 (rhTSG-6) injection does not significantly improve the delay of wound closure in diabetic HFD+STZ mice.</u> The rhTSG-6 protein (2 μg) was injected into each wound at Day 0, and at Day 2 post-wounding. The time course of reduction in wound area is depicted for four groups: CD + Vehicle, CD + TSG-6, HFD + STZ + Vehicle, and HFD + STZ + rhTSG-6. Statistical significance was assessed using the Mann-Whitney test.

### Supplementary Figure S5

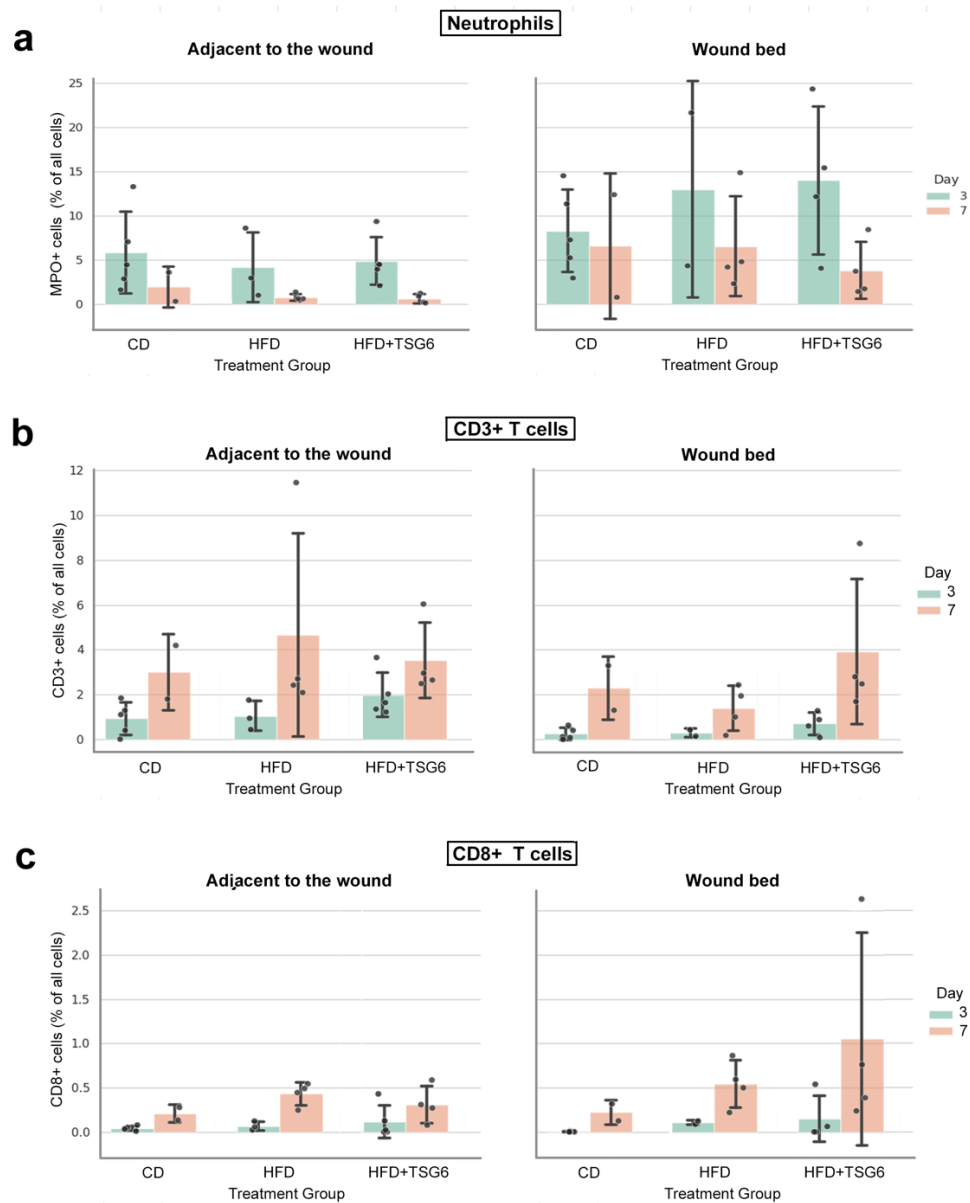

**Supplem Fig S5.** Recruitment of (a) neutrophils, (b) CD3+ (total T-cells), and (c) CD8+ (cytotoxic T-cells) in wounds at day 3 or day 7 post-wounding. Formalin-fixed tissue sections of wound biopsies were analyzed by multiplexed immunohistochemistry, wherein individual labelled cells were counted using QuPath software as described in **Supplemental Methods**. For each condition, each data point represents a different wound. Cell counts are shown from the central wound bed, and from the peri-wound skin immediately adjacent to the wound. The increases in the number of CD3+ and CD8+ T-cells at Day 7 versus Day 3, and decreases in neutrophils between Day 3 and 7, were as expected. However, no consistent differences between control diet-fed, HFD-fed, or HFD+STZ mice were observed for any of the three cell types shown.

### SUPPLEMENTARY MATERIALS AND METHODS

#### 1. Animals

C57BLKS/J ("wildtype" controls) and $Lepr^{db/db}$ (BKS.Cg-Dock7$+/+$Lepr db/J) male mice ("db/db" mutant mice) were purchased from Jackson Laboratories (JAX; Bar Harbor, ME) at six weeks old and wounded at eight weeks old after two weeks of acclimation. For the high fat diet + STZ experiments, C57BL/6J mice were purchased from Jackson Laboratories.

#### 2. Diets and treatments

For experiments to study the development of obesity and diabetes (including fasting blood glucose, glucose tolerance, and insulin resistance), 8-week-old C57BL/6J mice from JAX were assigned to one of three groups: HFD group, HFD+STZ group, or normoglycemic control, and started on either a high-fat diet (Cat. #D12331) or control diet (Cat. # D12329) from Research Diets, Inc., New Brunswick, NJ. The formulations of the two diets are shown in **Supp Table S1**. Streptozotocin (STZ) was obtained from Millipore-Sigma. At week five after diet initiation, the HFD+STZ group (alternatively called the "Diabetic" group in the manuscript) received daily intraperitoneal STZ injections (40 mg/kg) for three consecutive days, whereas the control group and HFD-only mice received vehicle injections (0.05 M citrate buffer). Mice were then maintained on their respective control or HFD until completion of the experiment. Such experiments included the following procedures (described separately below) in which blood was drawn from the tail vein at 7 days after the initial STZ injection and measured using a standard glucometer, to determine the following: (1) Fasting blood glucose levels (FBG); (2) Glucose tolerance test (GTT); (3) Insulin tolerance test (ITT). Wounding procedures were performed at approximately 10 days after the initial STZ injection. Mice were maintained per the American Association for the Accreditation of Laboratory Animal Care guidelines, and all procedures were approved by the Cleveland Clinic Institutional Animal Care and Use Committee (IACUC).

#### 3. Glucose tolerance test (GTT).

The mice were placed in a clean cage and subjected to a morning fast, during which they had access to only water for 6 hours (from 8:00 am to 2:00 pm). Following the fast, the mice were weighed. A glucose solution was prepared for intraperitoneal injection (1.5g per kilogram of body weight). The basal glucose levels were measured using a standard glucometer (Nova MAX PLUS, Nova Biomedical). Then, the glucose solution was injected via the intraperitoneal route (i.p.), and glucose levels were measured from the tail vein after 15, 30, 60, 90, and 120 minutes (Benede-Ubieto et al., 2020).

#### 4. Insulin resistance test (ITT).

Mice were placed in a clean cage with access to only water for a morning fast (6 hours; 8:00 am-2:00 pm). Mice were weighed, and an insulin solution was prepared for each mouse (0.75 IU insulin/kg body weight). If an animal became hypoglycemic with blood glucose below 20 mg/dL or appeared distressed, 300 µl of glucose solution (1.5 g/kg) was injected intraperitoneally (i.p.). Next, the basal level of glucose was measured using the glucometer. Then, insulin was injected via i.p., and glucose levels were measured in the tail vein blood after 15, 30, 60, 90, and 120 minutes.

### **5. Wounding protocols**

#### **5.1. Non-splinted wounding.**

Mice were anesthetized via intraperitoneal ketamine-xylazine injection. The upper back was shaved with an electric shaver. Weight and glucose measurements were done 24-48 hours before the day of wounding. Two full-thickness excisional wounds were made using 5-mm punch biopsies (Acuderm, Fort Lauderdale, FL). To monitor wound closure, mice were anesthetized with inhaled isoflurane and photographed using a digital camera on a stand with a fixed height. The wound area was determined from the digital images using NIH Image J software. In some experiments, skin and wounds were biopsied at pre-determined time points and proteins extracted as described under "Preparation of tissue lysates."

#### **5.2. Splinted wounding.**

In experiments using the db/db mice, wounds were created using <u>splints</u> as described by Wang et al. (Wang et al., 2013). Briefly, the dorsal surface of each mouse was shaved, and surface hair was removed with a brief 10-second application of Nair depilatory, followed by 3 minutes of rinsing with water. The next day, mice were anesthetized, and 5 mm punch biopsies were performed. A donut-shaped silicone stent with an outer diameter of 6 mm and an inner diameter of 5 mm was secured around the perimeter of the wound with the use of cyanoacrylate adhesive (Krazy glue, Elmer's product, Inc., Columbus, OH) plus eight interrupted sutures using 6-0 nylon sutures (6-0 Ethilon Nylon Suture, Ethicon L.L.C., Cornelia, GA). A trimmed transparent, bio-occlusive dressing (Opsite; Smith & Nephew, Inc) was placed on the splint. All splints were sterilized using a hydrogen peroxide system (Sterrad 100S system, Advanced Sterilization Products, Irvine, CA).

#### **5.3. Recombinant TSG-6 injections**

Immediately after wounding, recombinant human TSG-6 (rh-TSG-6, R&D Systems, Minneapolis, MN) in 100µl PBS was injected (total of 2 µg used for each wound). The 100 µl was injected at five sites in each wound, i.e., four sites at the edge plus the center of each wound. The source of rh-TSG-6 protein was a mouse myeloma cell line, NS0. Control wounds received PBS alone.

### **6. Preparation of tissue lysates**

On the day of harvest, mice were euthanized, and target tissues (skin wound, unwounded skin, visceral fat, subcutaneous fat, and muscle) were flash-frozen in liquid nitrogen and stored at -80°C for subsequent protein isolation. Frozen tissue samples were weighed and crushed using a tissue pulverizer on dry ice. Fresh lysis buffer (1M Tris-HCl @ pH 7.5, 5M NaCl, 0.05% Tween-20, and a 1:100 dilution of protease inhibitor cocktail) was added to the crushed tissue. The samples were homogenized on ice, vortexed every 2 minutes for 15 minutes, subjected to five quick pulses with an ultra-sonicator probe and centrifuged at 11,000 rpm for 6 minutes in a cold room. Supernatants were carefully transferred to a clean tube, and snap frozen on dry ice. The buffer used here was optimized for the multiplex cytokine assay (see below), but samples could also be used for Western blots or ELISA assays.

### **7. Western blots**

Lysates were subjected to electrophoresis on NuPage 4 -12% Bis-Tris gels (Invitrogen, Carlsbad, CA), blotted onto polyvinylidene fluoride membranes (Millipore, Burlington, MA), and probed overnight (4°C) with a target-specific primary antibody. The following antibodies were used:

• TSG-6 rabbit polyclonal (Abcam- ab204049)
• Actin(C-11)- rabbit polyclonal (Santa Cruz Biotechnology -Sc-1615 R)
• GAPDH(14C10)- rabbit monoclonal (Cell Signaling Technology-2118)

After removal of the primary antibody, membranes were washed and incubated with an appropriate horseradish peroxidase-conjugated secondary antibody, namely, goat anti-rabbit HRP (ROCKLAND, Gilbertville, PA # 611-1322) at 1:10000 concentration, at room temperature for 1.5 hours.

Chemiluminescent ECL Prime reagent kits (GE Healthcare, Chicago, IL) were used for detection. The blots were stripped and re-probed for actin or GAPDH for loading control. Protein bands were quantified using NIH Image J software.

### **8. Enzyme-linked immunosorbent assay (ELISA)**

According to the manufacturer's instructions, TSG-6 protein levels were quantified using the TSG-6 ELISA Kit (Mouse TNFAIP6 / TSG-6 ELISA Kit, LS-F7846-1, LSBio). Briefly, samples were added to microtiter plates pre-coated with an anti-TSG-6 antibody and incubated with a biotinylated detection antibody. Streptavidin-HRP was added, and the reaction was visualized using a TMB substrate. Optical densities were measured at 450 nm, and TSG-6 concentrations were determined by comparison to a standard curve.

### **9. Multiplex cytokine assay**

Cytokine levels in murine samples were quantified using the Mouse Cytokine Magnetic Kit (Catalog ID: MCYTOMAG-70K-07) from Millipore-Sigma, Inc, per the manufacturer's instructions. The principle of Multiplex Bead Array Assay (MBAA) is similar to ELISA, with the key distinction being the use of beads rather than a 96-well plate to hold the capture antibodies (Harris and Chen, 2019). In MBAA, capture antibodies are covalently attached to the surface of polystyrene beads. Each well in the assay can contain up to 100 bead types, with type of bead coupled with a different capture antibody. The bead conjugates are distinguished by the fluorescence intensity ratio of two or three different fluorescent dyes embedded within the bead. During sample incubation, the process is similar to that in ELISA, using biotinylated detection antibodies and a streptavidin-phycoerythrin (PE) conjugate reporter for detection.

Sample acquisition is performed using a specialized flow cytometer or fluorescent imager. This device executes a series of analyses similar to flow cytometric analysis of cell targets. Individual beads are first gated to remove doublets (beads that have adhered to one another) from the analysis. The bead type is then determined based on the ratio of the internal fluorescent dyes. Finally, the fluorescence intensity of the reporter is measured for each bead. This final measurement correlates with the concentration of a given analyte in solution, akin to the optical density readings in an ELISA assay. A standard curve is generated to calculate the final analyte concentration based on the median fluorescence intensity of the bead. Currently, two types of analyzers are used: MAGPIX® and FLEXMAP 3D®. The MAGPIX® analyzer uses a fluidics-based sample transport system and a magnetic plate to capture all beads simultaneously in a grid. Light-emitting diode (LED) lights illuminate the chamber, and a charge-coupled device (CCD) image sensor system captures a digital image of the beads to determine the fluorescence intensity of each bead. FLEXMAP 3D® combines three dyes within the microspheres, allowing for high-throughput analysis of up to 500 analytes and compatibility with 96- and 384-well plates; it uses three lasers to detect the bead regions and the reporter signal. We initially used the MAGPIX® system in our experiments and later transitioned to the FLEXMAP 3D® system because of its higher sensitivity.

### **10. Immunohistochemistry of skin wounds.**

Skin tissues were fixed in Histochoice (Amresco, Solon, OH), paraffin-embedded, and cut into 5-micron sections using a microtome. Hematoxylin and eosin (H&E) staining was performed for histological analysis of index sections. For analysis of macrophages, paraffin sections were rehydrated and blocked with 3% normal goat serum (30 min at room temp), exposed to primary antibody (Anti-F4/80 rat monoclonal, 1:100; Abcam), then to a biotinylated goat anti-rat biotin secondary antibody (1:300), followed by Vectastain ABC reagent (Vector Laboratories) and detection using a DAB peroxidase substrate kit. Slides were mounted, cover-slipped, and scanned using an Aperio AT2 (Leica Biosystems) digital pathology slide scanner.

For detection of other immune cells, sections were simultaneously immunostained for 3 antigens using the following primary antibodies: For neutrophils: (anti-Myeloperoxidase, 1:50; ab9535, Abcam); for CD3 T-cells: (anti-CD3, predilute; Roche); for CD8 T-cells: (anti-CD8, 1:100, Invitrogen). These were then followed by species-specific peroxidase-conjugated secondary antibodies and an individual Opal fluorophore (520 nm, 620 nm, or 690 nm) as provided by Akoya Biosciences (Marlborough, MA). Sample slides were scanned using the Vectra Phenolmager HT Automated Quantitative Pathology Imaging System, run by Phenolmager HT version 2.1.0 software (Quanterix, Billerica MA). Whole slides were scanned at 20x magnification using the appropriate dichroic filters and spectral unmixing library algorithm generated with Phenochart software v2.2.0 and Inform software v3.1.0 to spectrally unmix the scans.

### **11. Computer-assisted image analysis of inflammatory cell populations.**

#### **11.a. Quantification of Dermal Macrophages (F4/80)**

**METHOD:** Because macrophages in inflamed wound beds exhibit highly irregular and dendritic morphologies, standard nuclear-expansion cell segmentation algorithms are often error-prone. Therefore, we utilized a pixel classification algorithm written in QuPath (v0.7.0.; open-source, University of Edinburgh) (Bankhead et al., 2017) to quantify the overall F4/80 macrophage burden. Regions of interest (ROIs) were manually annotated to separate the wound bed (defined as intact tissue down to the panniculus carnosus) from the periwound region, explicitly excluding surface eschar and necrotic cellular debris. A pixel classifier was trained to identify DAB-positive (F4/80) pixels as positive signal and Hematoxylin-stained tissue as negative, while ignoring unstained areas. Data were quantified and presented as the percentage of F4/80-positive pixel area relative to the total cellular pixel area within the defined ROIs.

**RATIONALE FOR METHOD:** Macrophages often exhibit complex, dendritic morphologies in inflamed wound tissues. As a result, standard cell segmentation algorithms, which rely on uniform nuclear expansion, often fail to accurately capture these cells. Pixel classification overcomes this by quantifying the total area of positive target expression. Therefore, F4/80+ macrophages were quantified using a pixel classification approach in QuPath rather than cell-based segmentation. This method was selected because F4/80 is a membrane marker, and macrophages in inflamed wound tissue exhibit complex, dendritic morphologies with extensive cytoplasmic processes that preclude reliable individual cell segmentation. DAB and hematoxylin channels were separated using color deconvolution, and a trained pixel classifier was applied to quantify the percentage of DAB-positive area within annotated regions of interest.

#### **11.b. Quantification of Crown-Like Structures (CLS) in Adipose Tissue**

**METHOD:** High-fat diet (HFD) models induce adipocyte hypertrophy, triggering the recruitment of macrophages that encircle dead or dying adipocytes to form crown-like structures (CLS). Due to the variation in adipocyte size caused by hypertrophy, standard pixel classification was inconsistent for this compartment. To control for this, we normalized manual CLS counts to the total number of adipocytes per section. First, a QuPath pixel classifier was trained to identify the negative space of lipid droplets. These regions were refined using morphological filters (e.g., circularity and area) to create individual adipocyte objects. To determine average adipocyte size and total count per image, approximately 150 of these objects were randomly sampled per tissue section. CLS were then evaluated manually and defined as adipocytes exhibiting >50% encapsulation by F4/80-positive macrophages. Data were reported as the number of CLS per 1,000 adipocytes.

**RATIONALE FOR METHOD (CLS and Adipocyte Hypertrophy):** High-fat diets induce significant adipocyte hypertrophy. During this expansion, adipocytes can undergo apoptosis, triggering the recruitment of macrophages that arrange in crown-like structures (CLS) to clear the lipid debris. Because adipocyte size varies drastically due to hypertrophy, simple pixel classification of adipose tissue area is inconsistent and mathematically skewed. Normalizing manual CLS counts (>50% encapsulation) against the total number of adipocytes controls for this distortion, providing a more accurate measurement of macrophage presence within adipose tissue.

##### 11.c. Fluorescence Quantification of Neutrophils and T-cells

**METHOD:** For the assessment of neutrophils and T-cells, parallel sections were immunostained with antisera against MPO (neutrophils), CD3, and CD8 (T-cells) and imaged via confocal microscopy. The wound bed ROIs were defined in the same manner as the brightfield sections. Cell detection was performed within QuPath using the DAPI channel to identify cell nuclei, followed by a cell boundary expansion (3.5 $\mu\text{m}$) to capture cytoplasmic and membranous markers. To filter out non-specific autofluorescence and debris common in wound tissues, a Random Trees machine learning classifier was trained. The classifier evaluated cellular features to categorize the segmented cells into negative, MPO+, CD3+, or CD8+ populations based on their respective fluorophore intensities. Once visually validated for accuracy across samples, this optimized cell detection and classification algorithm was batch-applied to all images for consistency.

**RATIONALE FOR METHOD (Random Trees Machine Learning for Fluorescence):** Multiplex fluorescence is prone to background autofluorescence, especially in necrotic or highly fibrous regions of wound beds. Using a Random Trees ML classifier is superior to simple thresholding, as it evaluates a multidimensional set of cellular features (e.g. cytomorphology, staining intensity, texture) to distinguish signal (MPO+, CD3+, CD8+) from artifacts.

#### 13. Statistical analysis

For wound-healing experiments, a sample size estimate was performed using 80% power, a p-value 0.05, a 30% standard deviation, and literature on mouse models. With an expectation of observing a 50% difference, it was determined that at least six animals per group are needed. Data analysis was conducted using GraphPad Prism 9.0.0 software. Results are presented as means with standard errors of the mean (S.E.M.). The student's t-test was used to measure the data that passed four normality tests: Anderson-Darling, Kolmogorov-Smirnov, D'Agostino & Pearson, and Shapiro-Wilk. For all other data, the Mann-Whitney comparison was employed. A one-way ANOVA with a post hoc Tukey test was utilized to compare three or more groups. The threshold for statistical significance was set at $P < 0.05$, ensuring a robust interpretation of the results.
